# Trial-by-trial variability in cortical responses exhibits scaling in spatial correlations predicted from critical dynamics

**DOI:** 10.1101/2020.07.01.182014

**Authors:** Tiago L. Ribeiro, Shan Yu, Daniel A. Martin, Daniel Winkowski, Patrick Kanold, Dante R. Chialvo, Dietmar Plenz

## Abstract

Simple sensory stimuli or motor outputs engage large populations of neurons in the mammalian cortex. When stimuli or outputs repeat, the robust population response contrasts with fluctuating responses of individual neurons, known as trial-by-trial variability. To understand this apparent discrepancy, a detailed identification of the underlying spatiotemporal correlations is required. Here, we analyze spatial correlations in the instantaneous fluctuations between neurons relative to the neuronal population. Using 2-photon imaging of visual and auditory responses in primary cortices of awake mice, we show that these correlations grow linearly with the size of the observed cortical area. We extend these observations to the cortical mesoscale by analyzing local field potentials in behaving nonhuman primates. In network simulations, we demonstrate this linear growth in spatial correlation to emerge at criticality. Our findings suggest that trial-by-trial variability is a signature of critical dynamics in cortex maintaining robust, long-range spatial correlations among neurons.

## Introduction

Even simple stimuli or movements engage large numbers of neurons in the mammalian cortex. These robust population responses are contrasted by the heterogeneity and fluctuations of single neuron responses observed during repeated stimuli or motor outputs. This response variability or trial-by-trial variability has been consistently found *in vivo* for neurons that are non-selective to a particular stimulus feature (Deweese & Zador, 2004) as well as highly selective neurons (Heggelund & Albus, 1978; Tolhurst *et al*., 1983; Vogels *et al*., 1989; Shadlen & Newsome, 1998) and *in vitro* under network and stimulus conditions of reduced complexity (e.g. (Haroush & Marom, 2019)). While part of the variability has been attributed to single neuron properties, e.g. through a non-linear transfer between the membrane potential and firing rate (Carandini, 2004; Charles *et al*., 2018), numerous findings point to a significant role of the network dynamics to the observed variability. In particular, response variability is shared among neurons (Cohen & Kohn, 2011) through shared gain (Goris *et al*., 2014), includes the interactions between selective as well as non-selective neurons (Kotekal & MacLean, 2019), is supported by local cortical motifs such as fan-in (Dechery & MacLean, 2018) and depends on the balance of excitation and inhibition (Haroush & Marom, 2019). This potential role in network dynamics is in line with early findings that response variability strongly correlates with ongoing synaptic activity preceding a stimulus (Arieli *et al*., 1996).

The diversity and variability encountered in single neuron responses to even simple, repeated stimuli raises the question whether there exist certain principles that underlie this variability in the face of coherent network responses. These principles in order to be general should be independent from the specific stimulus given or motor output achieved, which translates into the general question of how individual elements, by interacting locally, can achieve a coherent, global population response. Correlation length measures can be used to identify the relationship between global, coherent system responses and fluctuating system components (Wilson, 1979). By studying the correlation in the fluctuations of system components, one can determine the distance at which they behave independently. It is well-established that the correlation length diverges at criticality (Wilson, 1976) in the thermodynamic limit. For finite systems, however, this asymptote behavior can be demonstrated by showing that the correlation length grows with system size. This relationship between correlation length and system size has been identified in the context of bird flocks, where a flock still maintains a coherent trajectory in space despite fluctuations in single bird trajectories (Cavagna *et al*., 2010; Bialek *et al*., 2012; Bialek *et al*., 2014). It was also demonstrated for resting activity of the human brain that the correlation length of the blood oxygenated level dependent (BOLD) signal scales linearly with the size of the brain region measured (Fraiman & Chialvo, 2012), a finding in line with expectations that the brain at rest maintains a critical state.

It has been argued (Chialvo, 2010) and shown experimentally (Shew & Plenz, 2013) that neuronal networks confer multiple benefits for being in the critical state, such as high dynamic range and maximal information capacity. For the bird flock, the critical state has been suggested to confer fast and coherent changes in flight direction in response to external perturbations, such as a predator attack (Bialek *et al*., 2014). For the brain, one might expect that external perturbations from sensory input or internal perturbations such as self-initiated motor commands lead to population responses that maintain the critical state. If this was correct, the correlation length for sensory and motor responses in the brain should scale with the size of the observed population of neurons and should be independent of feature selectivity. Here, we show experimentally a linear scaling in correlation length in the fluctuations of neuronal evoked responses in cortex. Using numerical simulations of a network model, we demonstrate this scaling to be found when the model is tuned to be critical. Our results identify a specific form of scaling in the trial-by-trial variability for sensory and motor responses in cortex *in vivo* and suggest this variability is a signature of critical dynamics that maintains robust, long-range correlations between neurons. Finally, by comparing experimental and numerical correlation functions, we are able to estimate the interaction length (i.e. the distance at which neurons interact physically) in visual cortex.

## Results

We investigated the extent in the spatial correlation of neuronal activity at the cellular and mesoscale level in cortex using two different experimental approaches. At cellular resolution, we studied pair-wise correlations between individual neurons using 2-photon imaging (2PI), whereas at the mesoscale in nonhuman primates we measured the local field potential (LFP) using high-density microelectrode arrays. We analyzed ongoing as well as activity obtained during sensory/motor processing for each scale. We will first present results obtained at the cellular level in awake mice followed by a comparison at the mesoscale in behaving nonhuman primates. Simulations of neuronal activity position our results in the context of subcritical, critical and supercritical dynamical regimes.

### Linear scaling of correlation lengths in mouse primary cortex

For cellular analysis, spiking activity in pyramidal cells of mouse primary visual cortex (V1; n = 7 mice) was recorded using 2PI over an area of *L x L* = ∼400 µm *x* 400 µm in superficial cortical layers at a depth of ∼100 – 150 µm from the cortical surface. Mice were quietly resting during recording while 2 s-lasting drifting gratings were presented every 4 s (Fig. 1a). As can be seen in the variance over average response plot (Fig.1b), the Fano factor was above one for most cells, indicating high response variability in single neuron responses. About 20% of neurons were tuned to stimulus direction with a selectivity index (DSI; Fig. 1c; see Materials and Methods) larger than 0.4 and an average response profile (Fig. 1d) depicting minimal response to the orthogonal orientation. Importantly, all single neuron responses were embedded in spatio-temporal clusters of activity that distributed in cluster size according to power laws, the hallmark of neuronal avalanches (Beggs & Plenz, 2003; Bellay *et al*., 2015). This scale-invariant organization in activity clusters was found during stimulation regardless of the directional drift as well as during luminance-matched gray screen presentation (Fig. 1e; *p* < 0.05 for power law test; see Materials and Methods).

**Fig. 1:**
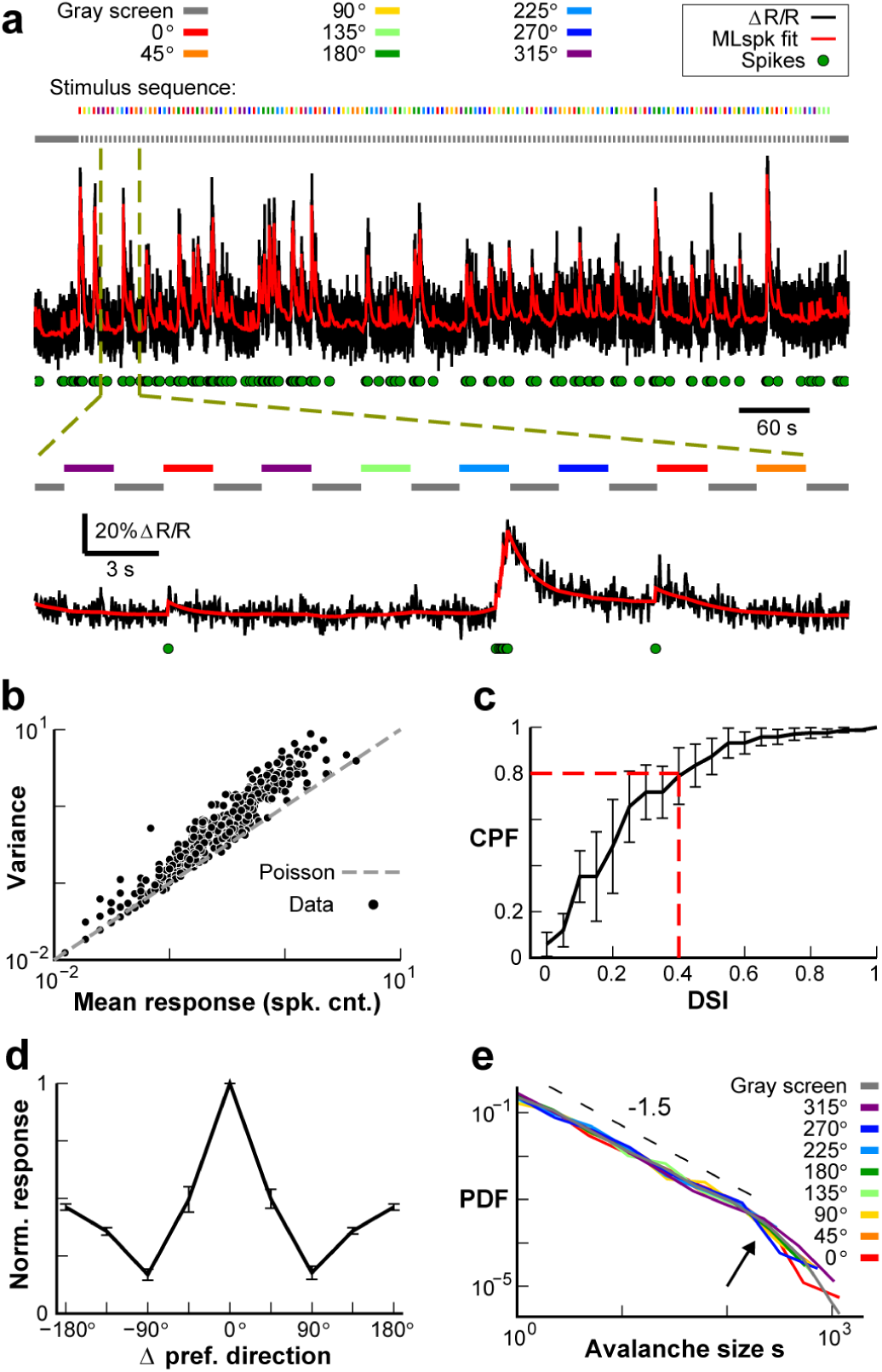
Trial-by-trial variability and tuning heterogeneity of single neurons during visually evoked avalanches in V1. **a** Variable response of a single V1 pyramidal neuron to semi-random presentation of 2 s large-field drifting gratings at 8 directions. *Black*: ΔR/R; *Red*: denoised ΔR/R; *Green circles*: estimated spikes (see Materials and Methods). *Color bars*: stimulus direction. *Gray bar*: stationary gray screen. *Broken lines*: Temporal zoom in for details. **b** Trial-by-trial variability of single neuron responses (*dots*; n = 228 cells across n = 7 animals), quantified by dividing response variance by response average, exceeds prediction from a Poisson process (*broken line*). **c** About 20% of responding neurons show a direction selective index (DSI) above 0.4. Cumulative probability function (CPF) for n = 7 animals (mean ± standard deviation). **d** Neurons with significant DSI show minimal response at orthogonal directions. Mean direction selectivity profile normalized to corresponding preferred direction (n = 228 cells; n = 7 animals; ± standard error). **e** Visually evoked responses in V1 organize as avalanches independent of stimulus direction, similarly to gray screen. Power law in avalanche size distributions (combined from n = 7 animals) with cut-off close to maximal number of ROIs (*arrow*). *Broken line*: slope of -1.5 as visual guide.

The response variability encountered in single neurons, their tuning diversity and scale-invariant grouping in the form of avalanches points to a fluctuation-dominated dynamic regime of cortex, in which individual elements seem to be difficult to relate to stable population responses. One potential answer to this problem could lie in critical dynamics, which maintain some order in the presence of strong fluctuations (Chialvo 2010). Therefore, to gain deeper insight into the dynamical regime of the cortex during the relatively short-lasting (compared to the scale of large avalanches) evoked responses and, at the same time, avoid the statistical influence of the strong drive created by the stimulus, we needed an alternative approach to the statistical distribution of observed activity patterns. Here we assessed the dynamics of the cortical network by analyzing how correlations in the evoked responses scale with the size of the observed system. We first directly calculated how the average pairwise correlation between neurons changes as a function of cortical distance *r* for the full recording window. This correlation was well described by a power law with exponent close to -1 and exponential cut-off, in line with expectations for critical dynamics (Cavagna *et al*., 2010) measured within a finite size window (Fig. 2b; *p* < 0.05 for power law test). Importantly, when activity from each of the trials for a given stimulus angle was randomly shuffled for each cell independently (trial shuffling; see Materials and Methods), the correlations dropped to zero regardless of the distance between the cells. This indicates that the total activity induced by the stimulus does not contribute to the distance related decay in correlations. We then assessed how the fluctuations of neuronal responses around the instantaneous population mean change with distance *r* between cells, by subtracting the population average as a function of time from each neuron’s time series (Fig. 2c). The correlation in fluctuations as a function of distance *r, C(r)*, was assessed for varying length *L* of the observation window (Fig. 2d). This variation in *L* systematically changes the population mean as exemplified in Fig. 2c for two window lengths. Indeed, nearby neurons deviate in similar fashion from population activity and *C(r)* decayed similarly, both for drifting gratings as well as during gray screen. At a certain distance, defined as the correlation length, ξ (Fig. 2d, arrows) (Cavagna *et al*., 2010; Fraiman & Chialvo, 2012), *C(r)* turns negative. We found that *C(r)* decayed similarly with distance for different *L* < 400 µm (Fig. 2d, different colors) and ξ was shorter for smaller windows.

**Fig. 2:**
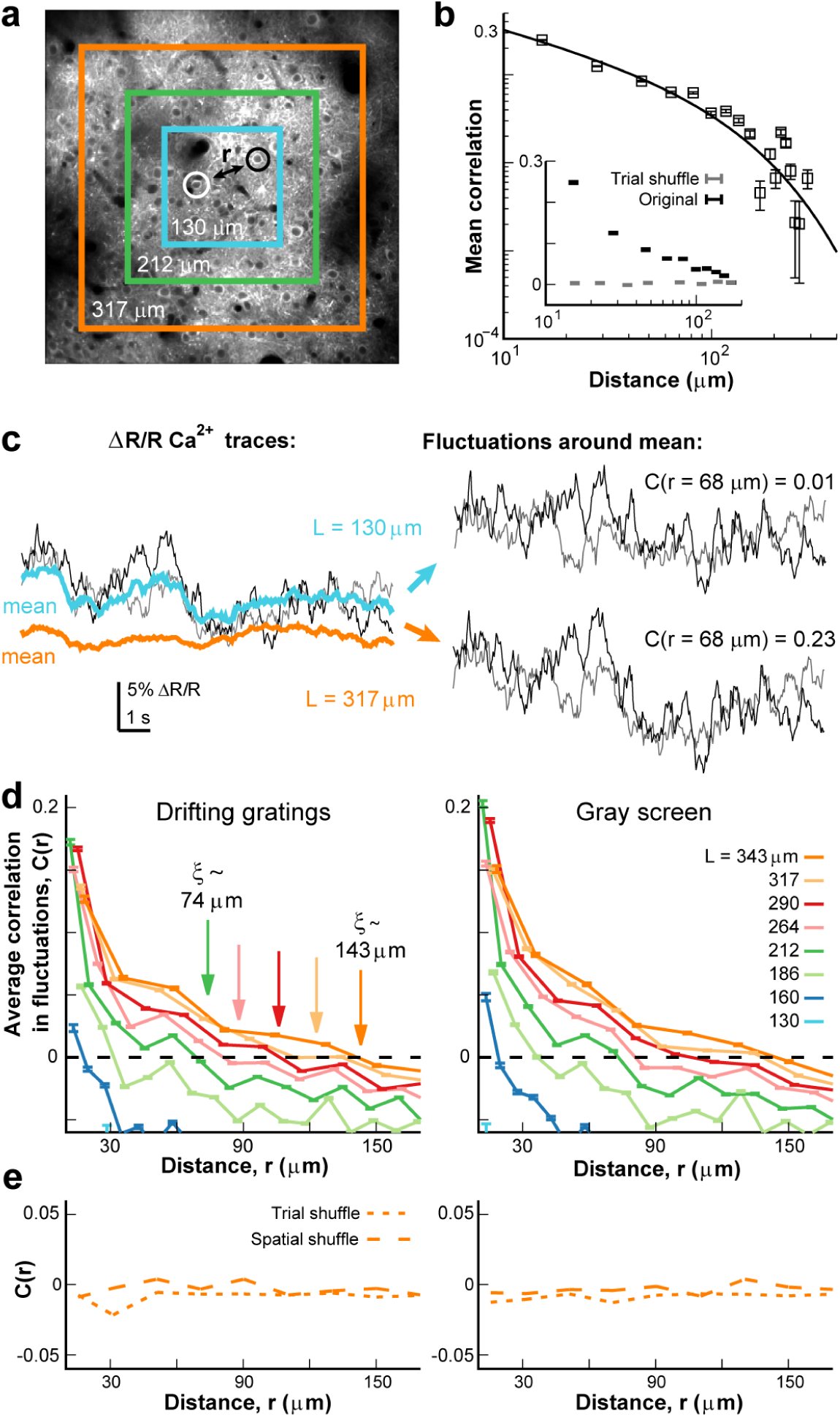
Correlations in activity fluctuations between V1 neurons grow with the size of the cortical area observed during visual presentations. **a** Schematic to measure pair-wise correlation between two V1 neurons (*open circles*) separated by distance *r* for different fields of view (*colored squares*), superimposed on a typical 400 x 400 μm 2-photon image showing YC2.6 expressing pyramidal neurons in V1 (∼120 μm cortical depth). **b** Average spatial correlation in ΔR/R for V1 neurons decays according to a power law with exponential decay during visual presentations in line with prediction for critical dynamics. *Squares*: data (mean ± standard deviation). *Line*: power law with exponential decay fit. *Inset*: Trial-shuffling (*gray*) results in near-zero correlations independent of distance. For comparison, original correlation (*black*) added in log-linear coordinates. *Bars* indicate standard deviation. **c** Example comparison of fluctuations in neuronal responses relative to the population response as a function of window extent. *Left*: Individual, single ΔR/R Ca^2+^ traces for two neurons (see panel **a**). Corresponding population activity fluctuates more in small windows (*blue*) compared to larger windows (*orange*; see panel **a**). *Right*: Subtracting the instantaneous population activity from individual neuronal responses reveals the corresponding fluctuations around the mean for each neuron as a function of window extent. Correlation value for each window extent is shown. **d** Correlation in the fluctuations around the mean decays with distance and crosses zero. The zero crossing defines the correlation length ξ, which is seen to increase with window length (*arrows*). Averages for n = 7 animals and all gratings directions (*left*) and gray screen (*right*). *Colors*: window length. *Broken line*: zero correlation. **e** Correlations are destroyed by trial-shuffling in time (*short dash*) or shuffling of ROI position (*long dash*), as revealed from the near-zero values obtained, independent of distance (*L* = 343 µm case shown).

Deviations from the population average of cells further apart than ξ were found, on average, to be anti-correlated for any window size. This anti-correlation is to be expected to a certain degree as by definition, at any given time some cells will exhibit activity above the population mean, while others will be below it. This fact and the general drop in correlations with distance will lead to a crossing through zero correlation, allowing to estimate ξ. We used trial shuffling as a control to demonstrate that ξ indeed captures emergent correlations in trial-by-trial fluctuations among neurons and does not reflect an artificial position due to subtracting the instantaneous mean. Trial shuffling maintains the common input introduced by the stimulus and distances among neurons, while removing coordinated changes of individual neuronal response around the population response. Trial shuffling results in *C(r)* being close to zero regardless of distance between neuron pairs, and therefore ξ cannot be estimated. This is true for both drifting gratings stimulus and gray-screen periods (Fig. 2e). Similarly, ξ is not defined after randomly permuting cell positions while keeping instant trial-by-trial correlations intact. These controls carried out for the full window size confirm ξ as a correlation length measure for neuronal activity that emerges from interactions between the observed neurons.

We note that a common way to obtain correlation length estimates from a system is precisely with the fit performed in Fig. 2b, by measuring how quickly correlations (without the population subtraction) drop with distance between system components after the power law regime. More specifically, the characteristic decay of the exponential portion of such a correlation spatial function can be used as a measure for the correlation length. However, as can be seen in Fig. 2b, the correlation function becomes very noisy for larger distances, reflecting the small number of pairs of neurons that far apart. Consequently, our estimates for the correlation length using that method become very unreliable (ξ ∼ 110.97 ± 58.55; 95% confidence interval). With the alternative method (Cavagna *et al*., 2010), which has previously been employed in neuronal activity data (Fraiman & Chialvo, 2012), we achieved a full collapse for all *L* by rescaling *C(r)* with its corresponding value ξ (Fig. 3a; *p* < 0.05; see Materials and Methods) and obtained a linear relationship for ξ over *L* (Fig. 3b), i.e. spatial correlations in V1 grow linearly with the observed cortical area. This increase holds regardless of sensory stimulus presented (Fig. 3b, different colors; *p* < 0.05 chi-square test for both cases; see Materials and Methods) and for subsets of cells responsive to the stimulus as well as those that are not (Fig. 3b, inset; *p* < 0.05 chi-square test for both cases), indicating that all cells, independent of their tuning-selectivity, follow the same correlation principles. Linear scaling with window size is also obtained when spike density estimates are being used (Suppl. Fig. S1). Our findings suggest that fluctuations around the mean between V1 neurons are scale-invariant, a hallmark of criticality.

**Fig. 3:**
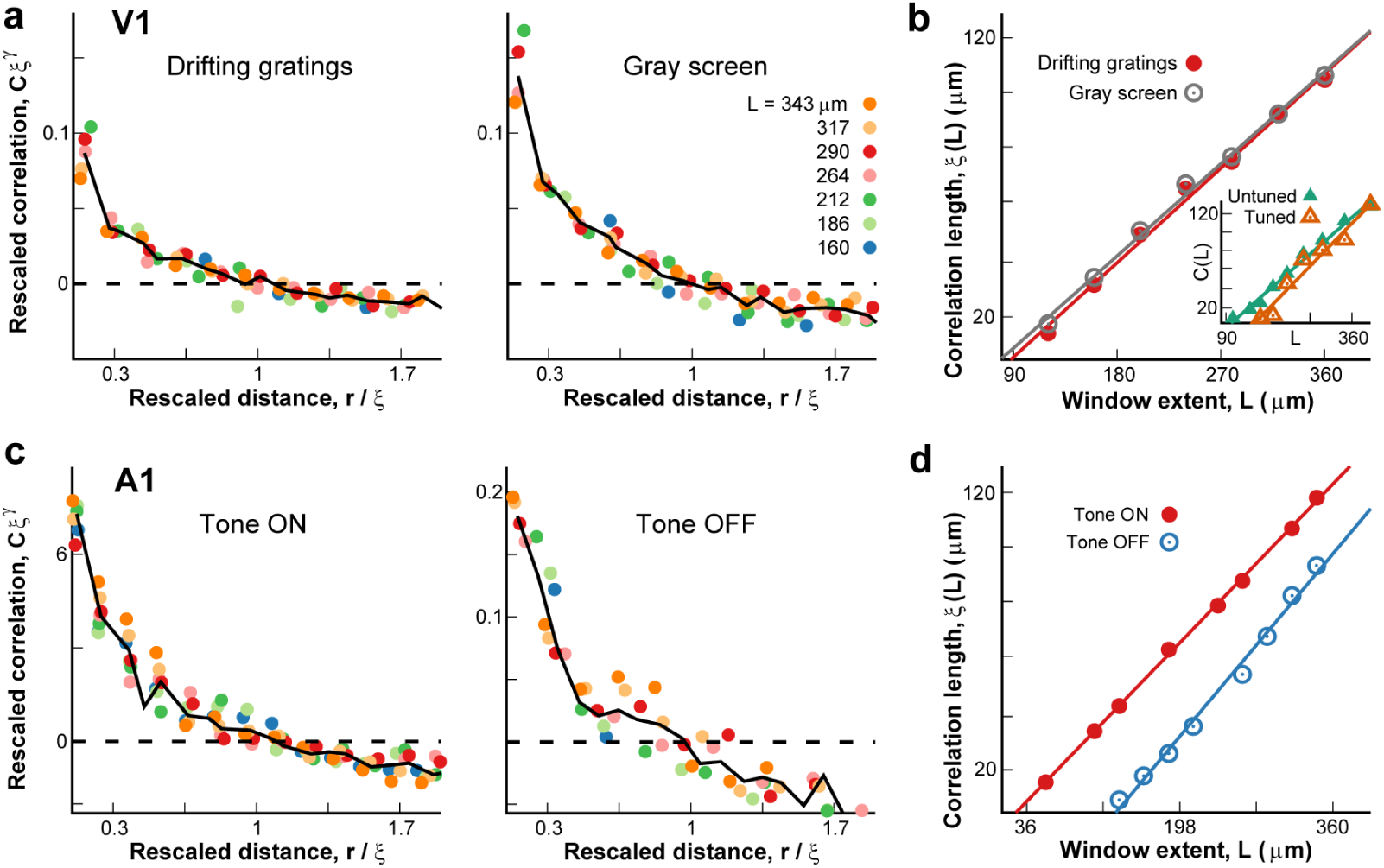
Linear growth in correlation length between neurons in primary visual and auditory cortex during sensory stimulation. **a** Scale-invariant spatial correlation function in V1 during sensory stimulation obtained by collapsing correlation functions from different window extents (cf. Fig. 2d; average from n = 7 mice). Rescaled correlations *C x* ξ^γ^ as function of distance normalized by correlation length ξ. *Black solid line*: average. *Left/right*: Drifting gratings/gray screen. **b** A linear growth in correlation length ξ with window extent *L* is found for drifting gratings (*red*) as well as gray screen (*gray*) presentations (n = 7 mice). *Inset*: tuned (*orange*) as well as untuned (*green*) neurons exhibit similar linear growth in spatial correlations. *Lines*: linear regression. **c** Neurons in primary auditory cortex of passively listening mice exhibit scale-invariant spatial correlation functions for both tone ON and OFF conditions (*left* and *right*, respectively). **d** Corresponding linear growth in correlation length ξ as function of *L*, for tone ON (*red*) and OFF (*blue*) (n = 11 mice). *Lines*: linear regression.

We extended these findings from V1 to the primary auditory cortex in awake mice. Mice passively listened to short (1 s) auditory tones semi-randomly presented from 8 different frequencies and 3 different volume levels every 3 – 5 s (Bowen *et al*., 2019). Evoked responses were recorded using 2PI in pyramidal neurons from superficial layers (n = 11 mice). Evoked neural responses were analyzed as described for visual cortex. Correlation functions *C(r)* were collapsed based on correlation length ξ and recording window size (Fig. 3c). In line with what was found for V1, ξ grew linearly with *L* for both tone ON and tone OFF conditions in auditory cortex (Fig. 3d; *p* < 0.05 chi-square test for both cases).

### Linear scaling of correlation lengths in nonhuman primates at cortical mesoscale

For the mesoscale level, i.e. analysis beyond several hundred µm, we employed microelectrode arrays (MEAs; 10 x 10 electrodes without corners; 400 µm interelectrode distance) and recorded the local field potential (LFP) in prefrontal (PF; monkey A) and premotor (PM; monkey B) cortex over an area of ∼4 mm x 4 mm (Fig. 4a; for details see (Yu *et al*., 2017)). Monkey A was trained in a visual-motor mapping task (Fig. 4b, bottom; see Materials and Methods), while monkey B performed a self-initiated movement task (Fig. 4b; top). We applied the methods described in the previous section to investigate whether the same results could be observed in this larger scale. Amplitude fluctuations in the LFP were obtained by subtracting the instantaneous average on the squared subarrays of width L from each electrode LFP. We then obtained *C(r)* between pairs of electrodes for the subarray of width *L* in multiples of the inter electrode distance (400 µm). *C(r)* decayed with distance (Fig. 4c) in remarkably similar shapes for different *L*, allowing for successful collapse of the curves (Fig. 4d). The collapsed functions were similar among both monkeys (Fig. 4d, left *vs*. right), during baseline or task-evoked activity (Fig. 4d, top *vs*. bottom). As found for primary visual and auditory cortex, ξ grew linearly with *L* for both cortical regions and was similar between baseline and motor/sensory processing epochs (Fig. 4e; *p* < 0.05 chi-square test for all cases). Importantly, this is not the case when trial shuffling is employed, as that procedure leads to a non-decaying near-zero correlation function (Fig. 4c, bottom), also in line with results obtained from mice.

**Fig. 4:**
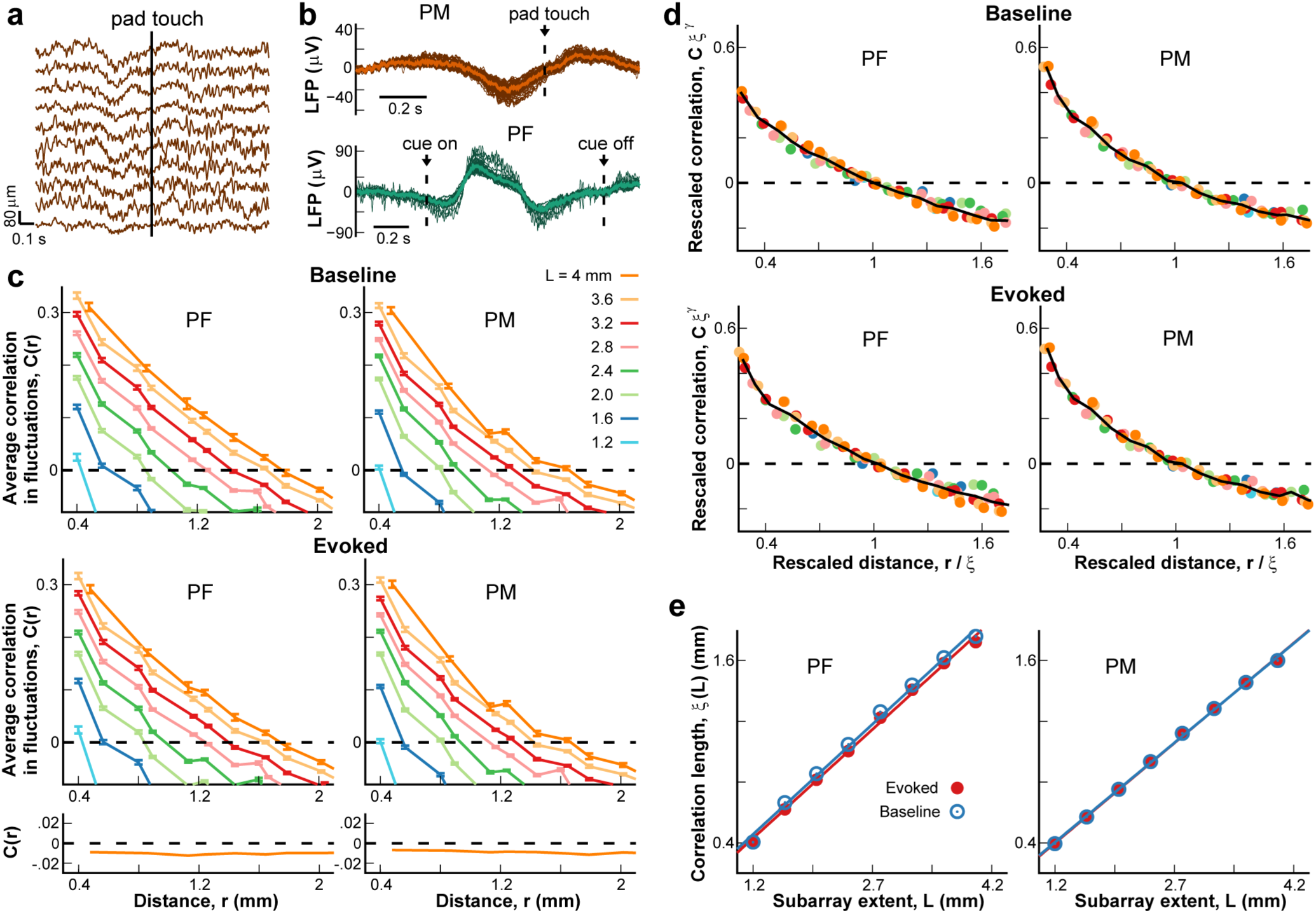
Scale-free correlation functions and corresponding linear growth in spatial correlations in monkey cortex during behavior. **a** Local field potential (LFP) traces during an example trial of a self-initiated motor task (10 example electrodes from a total of 96 electrodes at inter-electrode distance of 400 μm). **b** Trial-averaged LFP for the self-initiated motor task (*top*) to touch a pad (*broken line*; *arrow*) and a working memory task (*bottom*) around the presentation of a cue (*broken lines*; *arrow*). *Dark colors*: single electrodes. *Light trace*: population average. **c** Average correlation in fluctuations around the instantaneous population mean as function of distance for different subarrays of side length *L* (*color code*). *Top/bottom*: Baseline/task-evoked. *Left/right:* monkey A, PF/monkey B, PM. *Black dashed line*: zero correlation. *Bottom inset*: Near-zero correlations for trial-shuffled evoked datasets (*L* = 4 mm). **d** Rescaled correlations *C x* ξ^γ^ as function of distance normalized by correlation length ξ. *Black solid line*: average. **e** Correlation length growths linearly with subarray length *L* for monkey A PF (*left*) and monkey B PM (*right*) during baseline (*blue*) or task-evoked (*red*) activity. *Lines*: linear regression.

### Neuronal network model demonstrates linear growth of correlation lengths unique to critical dynamics

We simulated a neural network (see Materials and Methods and refs. (Kinouchi & Copelli, 2006; Ribeiro *et al*., 2014) for more details) to establish to which extent the results obtained in the experimental data can be used as an identifier for criticality. Specifically, we explore the impact of windowed access to a larger system, as this is currently the only way to examine finite-size effects on brain activity measures.

Furthermore, given the less than an order of magnitude spatial range we were able to access with 2PI and high-density microelectrode arrays, we estimated the expected deviation in correlation length estimates when networks move away from criticality. In the simulations we performed the correlation analysis exactly as described for the experimental data using a network of 1000 *x* 1000 units, with neurons interacting up to a distance *I*_*c*_ = 20. As can be seen in Fig. 5a, the correlations in the fluctuations were highest and extended for the longest distance in the critical regime (Fig. 5a, middle). Furthermore, the growth of ξ with the observed grid extent *L* was bounded for sub- and supercritical regimes (Fig. 5a, left and right), while in the critical regime, it increased with *L*. In fact, Fig. 5b shows that ξ peaked at criticality (σ ≃ 1; see Materials and Methods), regardless of the extent of the observed grid (Fig. 5b, different colors). We note that the absolute difference in the value of correlations from our simulations compared to experimental data simply results from the smoothing procedure employed in the latter (see Materials and Methods). In summary, the linear scaling of ξ as a function of *L* was unique to criticality, with sub- and supercritical systems presenting asymptotic behavior for ξ over *L* (Fig. 5c, different colors) and our simulations support our use of observation windows of different size as a proxy of system size in the analysis of correlations of critical systems. This approach was important so that the simulations properly mimic what is observed from the brain: only a small subset of the full network is then further subsampled in order to obtain the scaling of the correlation length. In Supplemental Figure S2, we provide further demonstration that this windowing approach (as opposed to measuring the correlation length as function of system size) is a valid method using the paradigmatic Ising model. Finally, in Fig. 5d we demonstrate that the susceptibility χ also peaks at criticality.

**Fig. 5:**
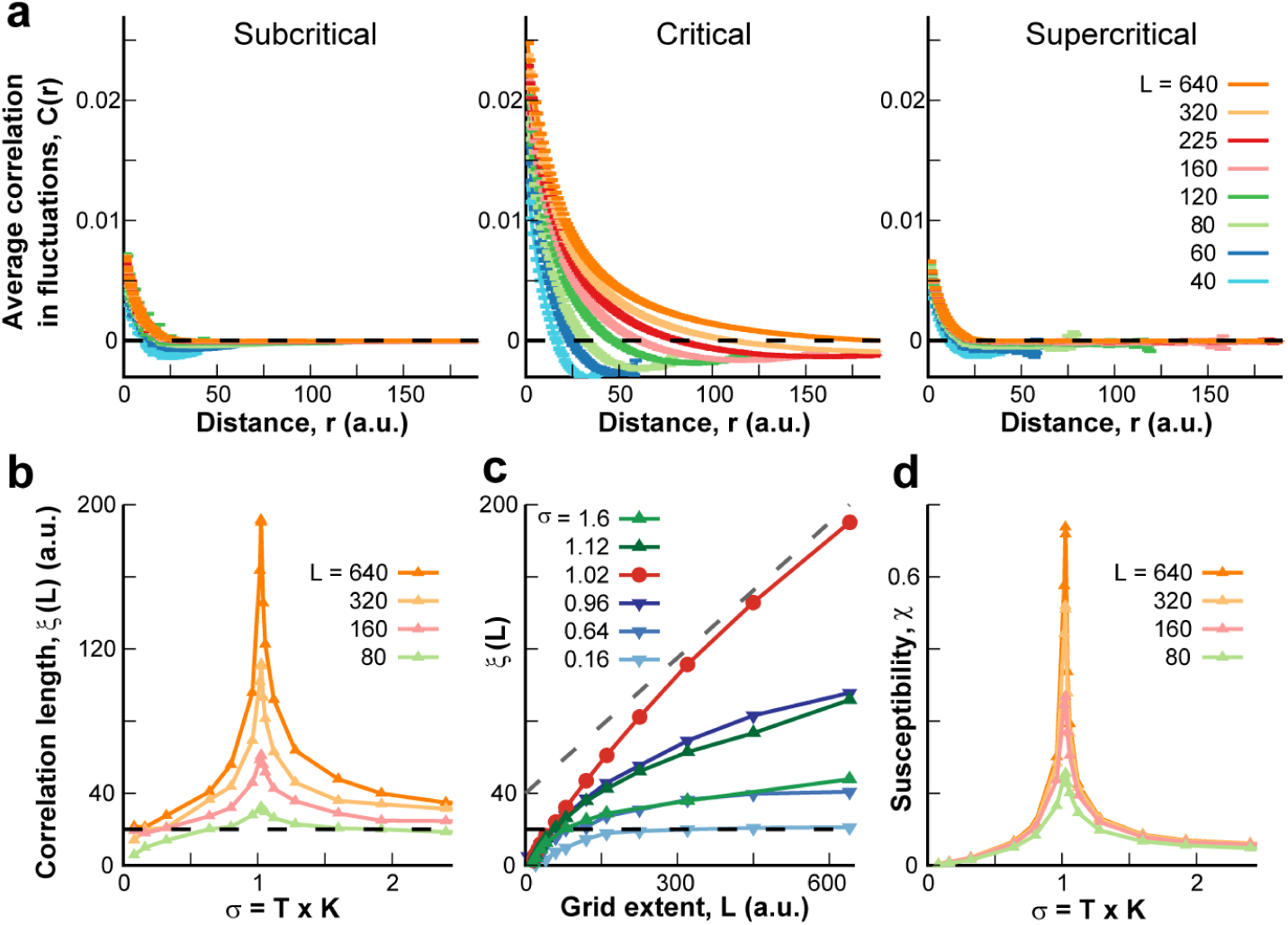
Neuronal model identifying linear growth in correlation length unique to critical dynamics. **a** Correlations in activity fluctuations as a function of distance for subcritical (σ = 0.64, *left*), critical (σ = 1.024, *middle*) and supercritical regime (σ = 2.4, *right*). *Color code*: the linear size of the observed window, or grid extent *L. Broken line*: zero correlation. *Insets*: Zoom-in to highlight the zero-crossings of the correlation functions. **b** Correlation length ξ as a function of σ (see Materials and Methods) for different grid extents (same *color code* as panel **a**). All curves peak for σ ∼ 1, corresponding to critical dynamics. **c** Correlation length ξ as a function of grid extent *L* for several values of σ (*color code*). *Dashed gray line*: linear growth with slope of 0.25 as a visual guide. Note the linear behavior of ξ for large *L* at criticality (σ = 1.02). Note also the subcritical case (σ = 0.16) for which ξ asymptotes at ∼20, which coincides with the interaction length *I*_*c*_ for the system (*dashed black line*). **d** Susceptibility X as a function of σ for different *L* (same *color code* as panel **a**). Note that all curves peak at σ ∼ 1, i.e. critical dynamics. All results in this figure were obtained from a network of size 1000 x 1000 with connectivity *K* = 16 and interaction length *I*_*c*_ = 20.

Altogether, these results demonstrate that at the critical regime, the range at which the fluctuations from two units in the network are correlated extends much further than the range at which units are connected, the interaction length *I*_*c*_ (see Materials and Methods). In contrast, for a system far away from the critical point, correlations cannot extend significantly beyond *I*_*c*_(Fig. 5c, dashed line at 20 coincides with asymptote value for ξ at σ = 0.16). However, as can be seen in Fig. 5c, even near the critical regime the linear growth of the correlation length is not present at smaller scales.

Finally, we studied how ξ(*L*) changed as a function of *I*_*c*_ for critical systems. Linear scaling was not obeyed for grid extents shorter than *I*_*c*_, for which the curve abruptly changed from the asymptotic slope to a sharper decay (Suppl. Fig. S3a, *left*; see also Supplementary Materials). Indeed, scaling ξ and *L* by *I*_*c*_, we obtained a collapse of all curves (Suppl. Fig. S3a, *right*). This scaling function clearly identified two regimes: scale free behavior took place for *L* > *I*_*c*_, while ξ increased much more abruptly for increasing *L* < *I*_*c*_, due to direct connections between units. We observed similar trends for individual mice in V1 when replotting the correlation lengths as a function of window extent with the linear regime being preceded by a steeper increase (Suppl. Fig. S3b, *left*). We estimated the point where the scaling abruptly changed for each mouse. They ranged from ∼100 to 180 µm for V1 networks. These estimates were used to successfully collapse all data (Suppl. Fig. S3, *right*).

## Discussion

Here, we used a correlation approach to assess the dynamical regime of the cortex during information processing epochs. The correlation length, a measure of how far apart neurons in cortex are positively correlated was shown to scale approximately linearly with the size of the observed window. If the distance over which neuronal activities correlate were to be finite, e.g. 200 μm, our approach would have revealed an upper bound in window size, or cortical area, for which the correlation length saturated. Instead, both of our experimental approaches, first, for up to 400 μm at the level of individual neuronal firing in primary visual and auditory cortex using 2-photon imaging and second, for up to 4 mm in the local field potential in awake nonhuman primates using high-density arrays, demonstrated that correlation length simply grows linearly with the size of the cortical area observed. Our simulations confirmed that our approach using windowing identifies linear growths in networks with critical dynamics, for which correlation length growths unbounded up to the final observed size.

We evaluated the correlation length as function of subsamples of the recorded region, i.e. compact windows, and not as a function of system size, which is the more common approach in physics. While it is known that windowing differs to a certain degree from finite size effects (Chen *et al*., 2011), we demonstrated for two different models that our method leads to the similar results. Given that the finite-size changes cannot be executed for real brains, windowing is therefore a realistic, alternative approach. Further support of our windowing method has been provided in BOLD fMRI recordings for the whole brain. In that study (Fraiman & Chialvo, 2012) it was shown that the linear growth in correlation length with the size of separate functional areas of the brain also describes the behavior of the correlation length when different subsets of multiple areas are added together in the analysis.

We used the fluctuations of local neuronal activity around the instantaneous mean of the observed network size to evaluate individual trials. The subtraction of the (observed) population average before calculating correlations has been successfully applied in the context of bird flocks (Cavagna *et al*., 2010), where one needs to evaluate how birds move in relation to one-another, disregarding the overall movement of the flock. This approach is much more common in physics (Wilson, 1976; Wilson, 1979) and, to our knowledge, has been applied here for the first time to single neurons and local neuronal populations in the wake animal. One might reconcile this common method by realizing that during sensory input or motor output, many neurons integrate similar inputs that consequently drive the activity of the network as a whole. Accordingly, subtracting the instantaneous population average reduces this ‘drive’ component, allowing the correlation analysis to focus on how activity in neurons changes in relation to one-another, independent from the general sensory/motor responses observed. This interpretation is thoroughly supported by our trial-shuffling results, in which the specific inter-neuron relationships that differ from trial-by-trial are removed and our correlation analysis correctly predicted zero correlations. We note that this method is not advisable for small number of neurons in which subtracting the average introduces dominant anti-correlations artificially decreasing correlation lengths for small windows. We mitigated this effect by imposing a minimum number of units in a given window before considering it for analysis. Small windows also suffer from the influence of the interaction length. Specifically, when the interaction length is on the order of the observed system size, the growth of the correlation length naturally is much faster with system size. We used this to our advantage, which allowed us to estimate the typical interaction range for superficial layers in the primary visual cortex.

That the cortical state is known to affect trial-by-trial variability (Kisley & Gerstein, 1999) and correlation among neurons (Cohen & Kohn, 2011; Rosenbaum *et al*., 2016) has been a long-standing observation. It is also well established that part of this variability originates from the cortical network itself (Kara *et al*., 2000; Sadagopan & Ferster, 2012; Goris *et al*., 2014; Schölvinck *et al*., 2015). Our demonstration of linear scaling in correlation length suggests critical dynamics as the framework that captures the intracortical correlation structure underlying this variability. Critical dynamics has been a fundamental driver in understanding optimization of information processing in complex systems in light of the evidence that fluctuations or variability are high at criticality (e.g. refs. (Shew *et al*., 2011; Fraiman & Chialvo, 2012; Tkačik *et al*., 2013; Karimipanah *et al*., 2017)). Decades ago, it was suggested that critical dynamics optimize information transfer in gene-regulation networks (Kauffman, 1969; Sole *et al*., 1999; Rämö *et al*., 2007; Nykter *et al*., 2008). Since then, the criticality hypothesis (Beggs & Plenz, 2003; Chialvo, 2010; Mora & Bialek, 2011; Plenz, 2012; Hesse & Gross, 2014; Marković & Gros, 2014; Plenz & Niebur, 2014; Bettinger, 2017; Cocchi *et al*., 2017; Muñoz, 2018) has gained much ground in the field of neuroscience. Highly desirable aspects of information processing have been shown to improve at criticality such as the maximization of mutual information between stimulus input and output (Kinouchi & Copelli, 2006; Shew *et al*., 2009; Shew & Plenz, 2013; Gautam *et al*., 2015; Bortolotto *et al*., 2016), increased information capacity (i.e. the number of possible internal states a network can establish) (Haldeman & Beggs, 2005; Shew *et al*., 2011; Tkačik *et al*., 2015), improved stimulus discrimination (Shriki & Yellin, 2016; Clawson *et al*., 2017), and the ability of neurons to flexibly change synchronization while maintaining an overall robust degree of phase-locking (Jantzen *et al*., 2009; Yang *et al*., 2012; Kelso *et al*., 2013; Kirst *et al*., 2017). Accordingly, our findings support the notion that trial-by-trial variability rather than reflecting pure noise, represents an intrinsic property of cortical dynamics during information processing.

Previous reports relied solely on the calculation of avalanche statistics in examining how close cortical networks are to critical dynamics. Such statistics is sensitive to non-stationarities in neuronal activity as found in evoked responses (Yu *et al*., 2017), which might explain the lack of avalanche statistics during sensory processing in *ex vivo* turtle (Shew *et al*., 2015) and the human MEG (Arviv *et al*., 2015). Power laws statistics as found for avalanches can in principle arise in critically balanced systems in the absence of correlations (see e.g. neutral avalanches (Martinello *et al*., 2017) and alternative models (Williams-García *et al*., 2014; Aitchison *et al*., 2016; Ioffe & Berry, 2017; Touboul & Destexhe, 2017). Our present results, therefore, which are based on the scaling of spatial correlations, provide a new experimental underpinning that cortical processing is in line with critical dynamics (Chialvo, 2010; Shew & Plenz, 2013).

## Materials and Methods

All procedures followed the Institute of Laboratory Animal Research (part of the National Research Council of the National Academy of Sciences) guidelines and were approved by the NIMH Animal Care and Use Committee or by the University of Maryland Institutional Animal Care and Use Committee.

### Mouse surgery and preparation

Wild type (C57/Bl6, Jackson Laboratory) mice were housed under a reversed 12 h-light/12 h-dark cycle with ad libitum access to food and water. Imaging experiments were generally performed near the end of the light and beginning of the dark cycle. A custom-made titanium head bar was surgically implanted onto the skull of the mice under isoflurane anesthesia (4% induction, 1-1.5% maintenance). A circular craniotomy (∼3 mm) was made above the area of interest (visual or auditory cortex), followed by injection of a virus containing the genetically encoded calcium indicator (YC2.6 for V1; GCaMP6s for A1) at a depth of ∼250-300 µm. After that, a cranial window composed of two 3 mm diameter coverslips glued to a 5 mm coverslip was implanted and the entire area (except for the window) was sealed with dental cement (Goldey *et al*., 2014; Bowen *et al*., 2019).

### Visual stimulation and response measures

Visual stimuli were prepared in Matlab using the Psychophysics Toolbox (Kleiner *et al*., 2007) and delivered via a monitor (Dell, 60 Hz refresh rate) placed ∼25 cm in front of the contra-lateral eye of the mouse. The stimulus was composed of moving gratings at 8 different directions presented for 2 s at maximum contrast, 0.04 cycles per degree and 2 cycles per sec. Stimuli were interspaced by gray screen (matched for average luminance) for 2 s. Each direction was presented 20 times in randomized order, for a total of 160 iterations. We calculated the direction selectivity index using the common definition: *DSI* = (*R*_*P*_ − *R*_*O*_)/(*R*_*P*_ + *R*_*O*_), where *R*_*p*_ and *R*_*o*_ are the responses to the preferred and opposite direction, respectively. Significance of *DSI* for each cell was assessed by comparing the values obtained from the original data with those obtained from shuffling the inter-spike intervals.

### Acoustic stimulation

Sound stimuli were synthesized in Matlab using custom software, passed through a multifunction processor (RX6, TDT), attenuated (PA5, Programmable Attenuator), and delivered via ES1 speaker placed ∼5 cm directly in front of the mouse. The sound system was calibrated between 2.5 and 80 kHz and showed a flat (± 3 dB) spectrum over this range. Overall SPL at 0 dB attenuation was ∼90 dB SPL on average (for tones). Sounds were played at a range of sound levels (40-80 dB SPL). Auditory stimuli consisted of sinusoidal amplitude-modulated (SAM) tones (20 Hz modulation, cosine phase), ranging from 3 to 48 kHz. The frequency resolution was 2 tones/octave (0.5 octave spacing). Each of these tonal stimuli was repeated 5 times with a 6 second inter-stimulus interval, for a total of 135 iterations (Bowen *et al*., 2019).

### Two-photon imaging and analysis

Images were acquired by a scanning microscope (Bergamo II series, B248, Thorlabs) coupled to a pulsed femtosecond Ti:Sapphire 2-photon laser with dispersion compensation (Vision S, Coherent). The microscope was controlled by ThorImageLS software. The wavelength was tuned to either 830 nm or 940 nm in order to excite YC2.6 or GCaMP6s, respectively. Signals were collected through a 16× 0.8 NA microscope objective (Nikon). Emitted photons were directed through 525/50 nm (green) and 607/70 nm (red) band filters (for GCaMP6s) or 535/22 nm (yellow) and 479/40 nm (cyan) band filters (for YC2.6) onto GaAsP photomultiplier tubes. The field of view was ∼400 x 400 μm. Imaging frames of 512×512 pixels were acquired at 30 Hz by bidirectional scanning of an 8 kHz resonant scanner. Beam turnarounds at the edges of the image were blanked with a Pockels cell. The average power for imaging was <70 mW, measured at the sample. The obtained images were corrected for motion using dft registration software with Matlab (Guizar-Sicairos *et al*., 2008). Regions of interest (ROI) were identified from the average image of the motion corrected sequence using custom code. For each labeled neuron, raw fluorescence signals (cyan and yellow for YC2.6; green for GCaMP6s) over time were extracted from the ROI overlying the soma. The mean ratiometric signal (R; YC2.6) or single fluorescence (F; GCaMP6s) in each ROI was calculated across frames and converted to a relative fluorescence measure (ΔR/R_0_ or ΔF/F_0_). The baseline signal R_0_ (or F_0_) was estimated by using a sliding window that calculated the average fluorescence of points less than the 10^th^-percentile during the previous 1.3-second window (40 frames).

### Monkey behavioral training and electrophysiological setup

Experiments were described previously (Yu *et al*., 2017). In short, two adult rhesus monkeys (*Macaca mulatta*) were surgically implanted with a titanium head post. After recovery, they were trained to sit head-fixed in a primate chair for behavioral performance. In the cue-initiated task, monkey A (male, 9 years old, 8 kg) had to press a bar in front of the chair upon presentation of the ‘trial-initiation’ cue on a computer screen. After ∼2 s, the initiation cue was followed by an ‘instruction’ cue, for the duration of 1 s. Upon cue disappearance, monkey A had to release the bar and reach with his right arm to one of two specialized feeders, depending on which of two possible cues were presented (Mitz *et al*., 2001). Approaching the incorrect feeder rapidly triggered a proximity sensor to sequester the food rewards in both feeders, which prevented the monkey from obtaining a reward on that trial. The inter trial interval was 3 – 5 s. In the self-initiated motor task, monkey B (female, 8 years old, 7 kg) had to move her right arm to touch a pad placed ∼30 cm in front of the monkey chair after which a food reward was given. After the monkeys learned their respective tasks, a multi-electrode array (MEA; 96 channels - 10×10 without corners, inter-electrode distance: 400 μm; electrode length: 1 mm for monkey A and 0.55 mm for monkey B; BlackRock Microsystems) was chronically implanted in the arm representative region of the left prefrontal area (area 46, monkey A) or the left premotor cortex (monkey B). The LFP (1 – 100 Hz band pass filtered; 2 kHz sampling frequency) was obtained from the implanted MEA. Electrophysiological signals as well as the timing of behaviorally relevant events, e.g. touching the pad, presentation of visual cues, etc., were stored for off-line analysis.

### Numerical simulations

We simulated a neural network, as described previously (Kinouchi & Copelli, 2006; Ribeiro *et al*., 2014). In short, each neuron can be in one of three states at each time step: 0 for resting, 1 for active, and 2 for refractory. The model considers *S*^2^ neurons on a square lattice. Each neuron outputs to *K* other neurons, selected with an exponentially decaying probability function of the Euclidian distance *r* between them (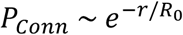, with R_0_= 5). A spatial cutoff is set in the interaction distance: neurons cannot directly connect at distances greater than *I*_*c*_ = 4*R*_0_ spatial units (thus, defining an interaction length *I*_*c*_). Furthermore, to reduce small *S* effects, we employed periodic boundary conditions. Results were computed on a square grid of length *L*. Simulations parameters were always such that *S* was much larger than *L* to reduce finite-size artifacts and to better mimic experimental data. A small Poisson drive (*h* = 10^−7^ per time step) to each neuron determined the overall rate of firing activity. The present results were robust over a wide range of *h* values (e.g., *h* = 10^−9^ to 10^−4^ per time step). The control parameter of the model determines the branching of the neural activity and was defined as σ = *K* × *T*, where *T* is the probability that an active neuron (i.e., in state 1) can excite each one of the *K* neighbors that it connects to. Therefore, as shown previously, the model can be made critical by selecting a transmission probability *T* such that σ ∼ 1, for any given *K*. Susceptibility was defined as the area under the correlation *vs*. distance curve, for a given window size observed, up to the correlation length.

Considering the focus of the present study, we note that there are four length scales in the model. The *interaction length* (here called *I*_*c*_) is the scale at which neurons can interact via direct connections. *System length* (called *S*) determines system size (the network is composed of *S*^2^ neurons). The third one is the window length *L* (*L* ≤ S), called *grid length*, which determines how many neurons we will measure from (i.e., *L*^2^ neurons). The last scale is the *correlation length* ξ, the longest distance at which on average the activity of any two given neurons may remain positively correlated. It is calculated from fluctuations on activity, following Cavagna’s work (Cavagna *et al*., 2010) (see section below). To avoid confusions, we remark that the term *interaction* is reserved here to denote *direct connections* and *correlations* to the mathematical result from computing *correlations of neural activity*. Simulations for varying interaction length (*I*_*c*_) were carried out using the same model described above. When changing *I*_*c*_, the decaying parameter used for making connections *R*_*0*_ was changed accordingly (such that the relation *T*_*c*_ = 4*R*_0_ still holds).

### Avalanche analysis

Avalanche analysis for the 2PI data was performed as described previously (Beggs & Plenz, 2003; Bellay *et al*., 2015). In short, estimated spikes (Deneux *et al*., 2016) from all neurons were pooled together to create a population activity series. Avalanches are defined by contiguous non-zero population activity preceded and followed by blank frames (frames with zero population activity). Avalanche sizes were defined by the total number of spikes throughout their lifetime. Power-law goodness of fit was evaluated through a p-value calculated from the log-likelihood ratio when comparing power law, exponential and lognormal fits, as described previously (Clauset *et al*., 2009; Klaus *et al*., 2011; Bellay *et al*., 2015).

### Correlation analysis

The correlation of the fluctuations as function of distance (Cavagna *et al*., 2010) was calculated as

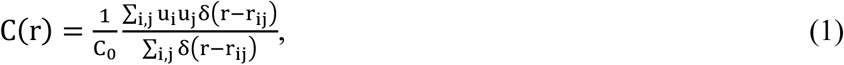

where δ(*r – r*_*ij*_) is a smoothed Dirac δ function defining all pairs of neurons located at mutual distance *r, r*_*ij*_is the Euclidean distance from the *i-*th neuron’s spatial location to the spatial location of neuron *j*, and *u*_*i*_ is the value of the signal γ of neuron *i* at time *t*, after subtracting the overall mean of signals γ from neurons inside the observation window of size *L* at that time 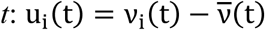.To ensure that *C*(*r =0*) = 1, the normalization factor 1/*C*_*0*_was used. We note that since the instantaneous average is subtracted,*C*(*r*) is not equivalent to the most commonly used pairwise Pearson correlation function.

The objective of computing the *C*(*r*) is to determine the correlation length ξ, which is defined as the point where the correlations of the fluctuations reaches zero, i.e. *C*(ξ) = 0. Since system size, i.e. cortex size in our experimental data, was fixed, we investigated how ξ changes with system size subsampled by our recordings and considered neurons/electrodes within a window of length *L* (this proxy is validated by computing correlation lengths both as function of increasing system and window sizes in a model (Suppl. Fig. S2). More specifically, for the 2PI data in mice, fields of view ranging from ∼ 40 × 40 μm (windows with fewer than 5 units were ignored to avoid bias introduced by the average subtraction procedure when the number of units is too small) to the maximum possible size were considered, while for the monkey LFP data the smallest subarray considered was 3 × 3. To reduce noise effects, results were averaged across all possible subregions for any given size. The time series were smoothed in the time domain (using Matlab routine *medfilt1*.*m* with 20 samples for the mice 2PI data, 8 samples for the monkey LFP data). This smoothing procedure improved statistics without changing the results qualitatively. In order to more precisely estimate the zero-crossing point for the experimental data, we fit 3^rd^ order polynomial functions to the *C*(*r*) curves around the zero-crossing.

To quantify linear growth in correlation length ξ as function of window length *L*, we first obtained a linear regression of the ξ(*L*) data followed by chi-square statistics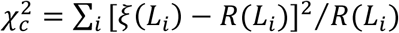, where *L*_*i*_ is the *i*^*th*^ measured value of *L* and *R*(*L*_*i*_) is the linear regression value at *L*_*i*_. 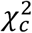 can be used to obtain a p-value that estimates how likely the data fit the linear regression that well by chance from the chi-square distribution. We rescaled the correlation *vs*. distance curves by normalizing the distances by the correlation length and by rescaling the correlations by the correlation length to the power of γ, defined as the slope of the curves at zero-crossing (Cavagna *et al*., 2010).

### Trial shuffling and spatial shuffling

Trial shuffling for the V1 data was obtained by randomly permuting the responses from each of the 8 presented directions separately. This was done for each neuron independently. Therefore, in each trial of the trial shuffled dataset activity from each cell corresponds to a response to the same stimulus presented in the original data but taken from different presentations of that stimulus. For example, suppose we presented stimulus 5 in trial 1. In the shuffled data, the response of neuron 1 in that trial may be taken from the 10^th^ presentation of stimulus 5, while response of neuron 2 may be taken from the 3^rd^ presentation. Spatial shuffling was performed by randomly permuting cell positions, leaving everything else unchanged.

## Acknowledgements

We thank members of the Plenz lab for lively discussions. This research was supported by the Division of the Intramural Research Program (DIRP) of the National Institute of Mental Health (NIMH), USA, ZIA MH00297 and the BRAIN initiative Grant U19 NS107464-01. D.A.M. acknowledges financial support from ANPCyT Grant No. PICT-2016-3874 (Argentina). This research utilized the computational resources of Biowulf (http://hpc.nih.gov) at the National Institutes of Health (NIH), USA, and UnCaFiQT-INIFTA (SNCAD), Argentina.

## Contributions

T.L.R., D.R.C. and D.P. conceived and planned the study; T.L.R., S.Y., D.W., P.K. and D.P. oversaw and carried out experiments; T.L.R., D.A.M., D.R.C and D.P. designed the models and carried out the simulations; T.L.R. took the lead in the analysis; T.L.R., D.R.C. and D.P. wrote the paper.

## Conflicts of Interest

The authors declare no competing financial interests.

## Supplementary Material

### Scaling of correlation lengths in mouse V1 based on spike density estimates

**Supplemental Figure S1:**
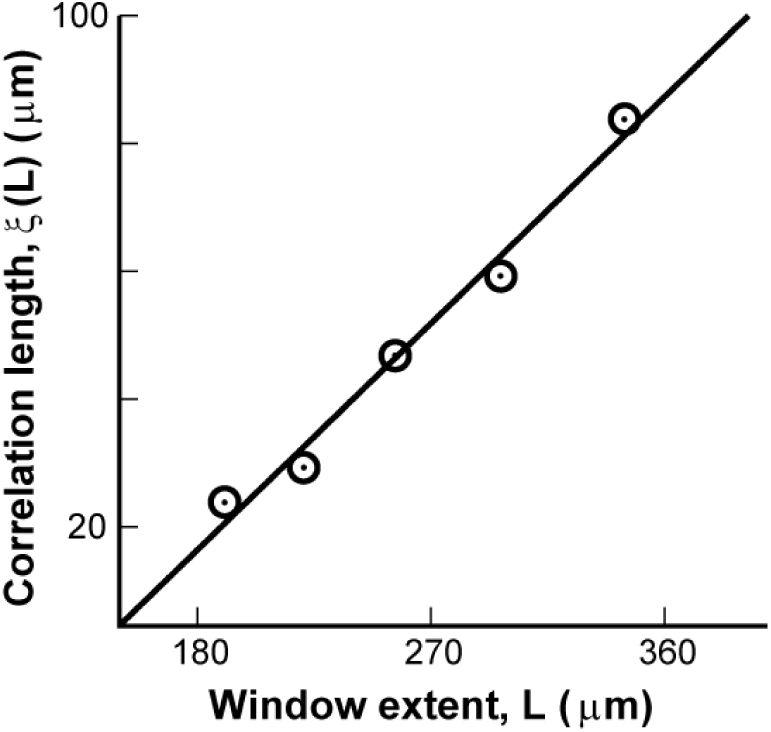
Correlation length scales linearly with window extent for spiking data. Correlation as a function of window extent obtained from estimated spike times from V1 in mice. *Line* represents a linear regression.

### Correlation length scaling for increasing window size in the Ising model

In critical systems, the correlation length is known to scale with system size. Because in biological systems changing system size is often impractical, in this work we introduce a proxy which uses windows of different sizes. In order to validate that approach, we test the concept in the paradigmatic 2D Ferromagnetic Ising model, which undergoes a critical transition at temperature T ≃ 2.3. The simulations (lasting at least 10^5^ Montecarlo steps) used two setups: in the first, the correlation function was computed in the standard way, from a model running on square lattices of increasing *S* = 16, 32, 64 and 128. In the second setup, a relatively large *S* = 300 square lattice (i.e. 300 × 300 spins) was simulated, and the correlation function was computed from square windows of smaller sizes *L* = 4, 8, 16, 32, 64 and 128. Representative results for three different temperatures are shown: subcritical (T = 1.6, Fig. S2a), critical (T = 2.3, Fig. S2b) and supercritical (T = 3.0, Fig. S2c). Fig. S2d shows the correlation lengths computed from both setups are very similar. Therefore, the curves obtained changing system size *S* or changing window size *L* produce similar correlation lengths. These results validate the use of window size as a proxy of system size in the analysis of correlations of critical systems.

**Supplemental Figure S2:**
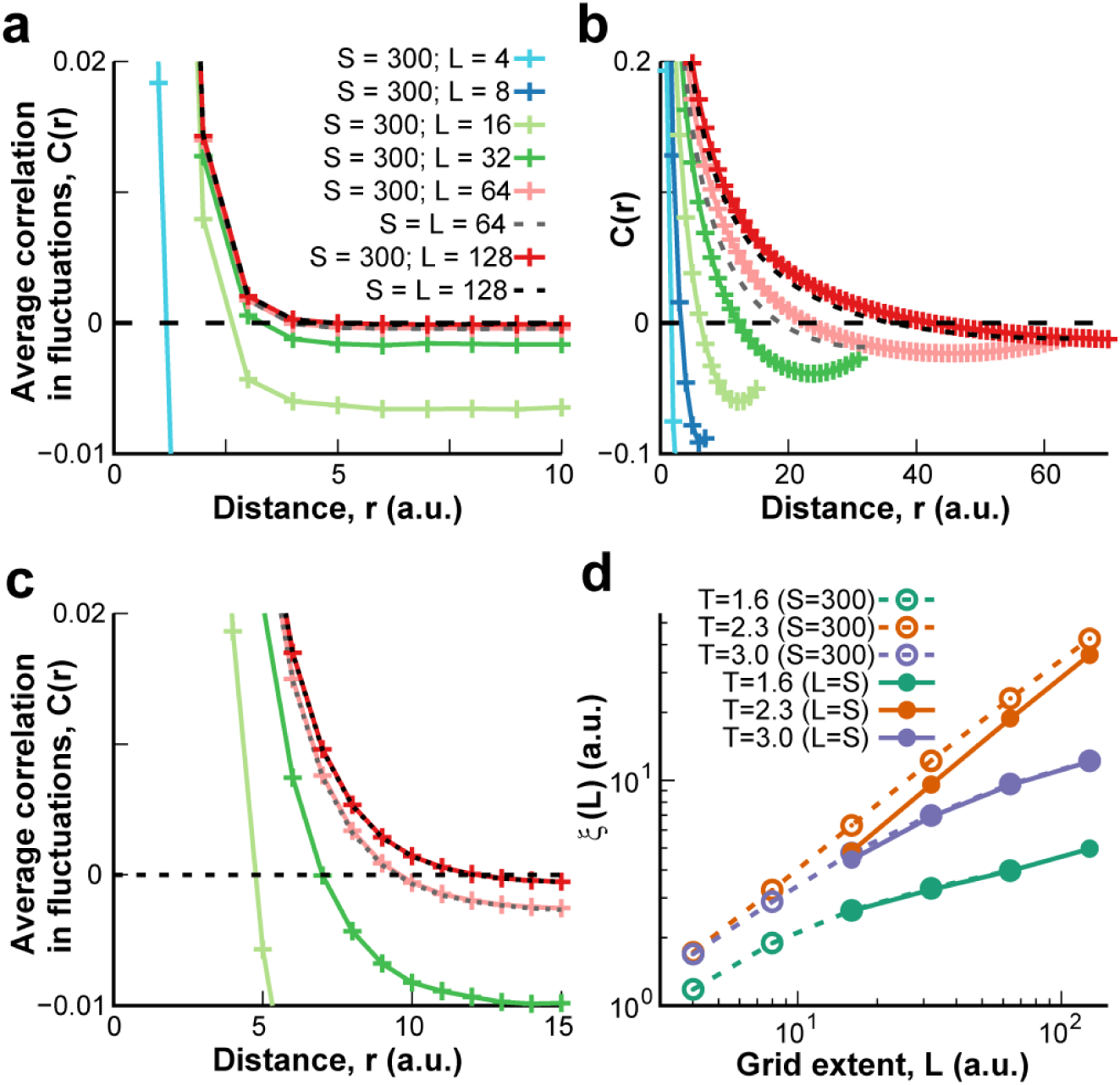
The ferromagnetic 2D Ising model. **a** Correlation as a function of distance for the subcritical regime (*T* = 1.6). *Color code*: different window sizes (*L* from 4 to 128) in a system of size *S* = 300. *Dotted lines*: fixed *S* = *L* = 64 and 128, as indicated. **b** Results for the critical regime (*T* = 2.3). **c** Results for supercritical regime (*T* = 3.0). **d** Correlation length as a function of distance for *T* = 1.6, *T* = 2.3 and *T* = 3 and different conditions. *Empty circles* and *dashed lines* are used for systems with *S* = 300. *Filled symbols* with *continuous lines* are used for systems and windows of sizes *S* = *L*.

### Estimating the interaction length from experimental data

We studied how the scaling law changed as a function of the interaction length for critical systems. We fit a power law of exponent 1 with an extended exponential part to the data as follows ξ(*L*) = *L*{1 - exp[-(*L*/*L*_0_)*L*^ɛ^]}, where *C* = 0.27 is a constant that controls the slope of the curves (in linear coordinates), ɛ = 3 is a constant that controls how sharp the decay for small *L* is and *L*_*0*_ is the cross-over point where the behavior changes from the exponential to the linear growth, our estimate for the interaction length. The values for *C* and ɛ were found empirically and represent a good match for the curves obtained in the simulations.

As shown in Fig. S3a (left), the linear scaling was not obeyed for grid extents shorter than *I*_*c*_, for which the curve abruptly changed from the asymptotic slope to a sharper decay. If this change in behavior was solely due to the influence of direct connections between units (as opposed to influence of critical dynamics) then we should be able to obtain a universal curve that is invariant to the range at which units can be connected. Indeed, scaling ξ and *L* by *I*_*c*_ we obtained a collapse of all curves (see Fig. S3a, right). This scaling function clearly identified two regimes: scale free behavior took place for *L* > *I*_*c*_, while ξ increased much more abruptly for increasing *L* < *I*_*c*_, due to direct connections between units. We observed similar trends for individual mice in V1 when replotting the correlation lengths as a function of window extent with the linear regime being preceeded by a steeper increase (Fig. S3b, left). The point where the scaling abruptly changed differed between mice (Fig. S3b, different colors) and we used a funtion that mimics the behavior observed for the collapsed curve in Fig. S3a (right) to estimate the interaction length for each mouse (see Materials and Methods). These estimates were used to collapse all functions (Fig. S3b, right). They ranged from ∼100 to 180 µm among mice and are a good estimate for the local distance at which pyramidal cells typically connect in superficial layers in V1 (Levy & Reyes, 2012; Seeman *et al*., 2018).

**Supplemental Figure S3:**
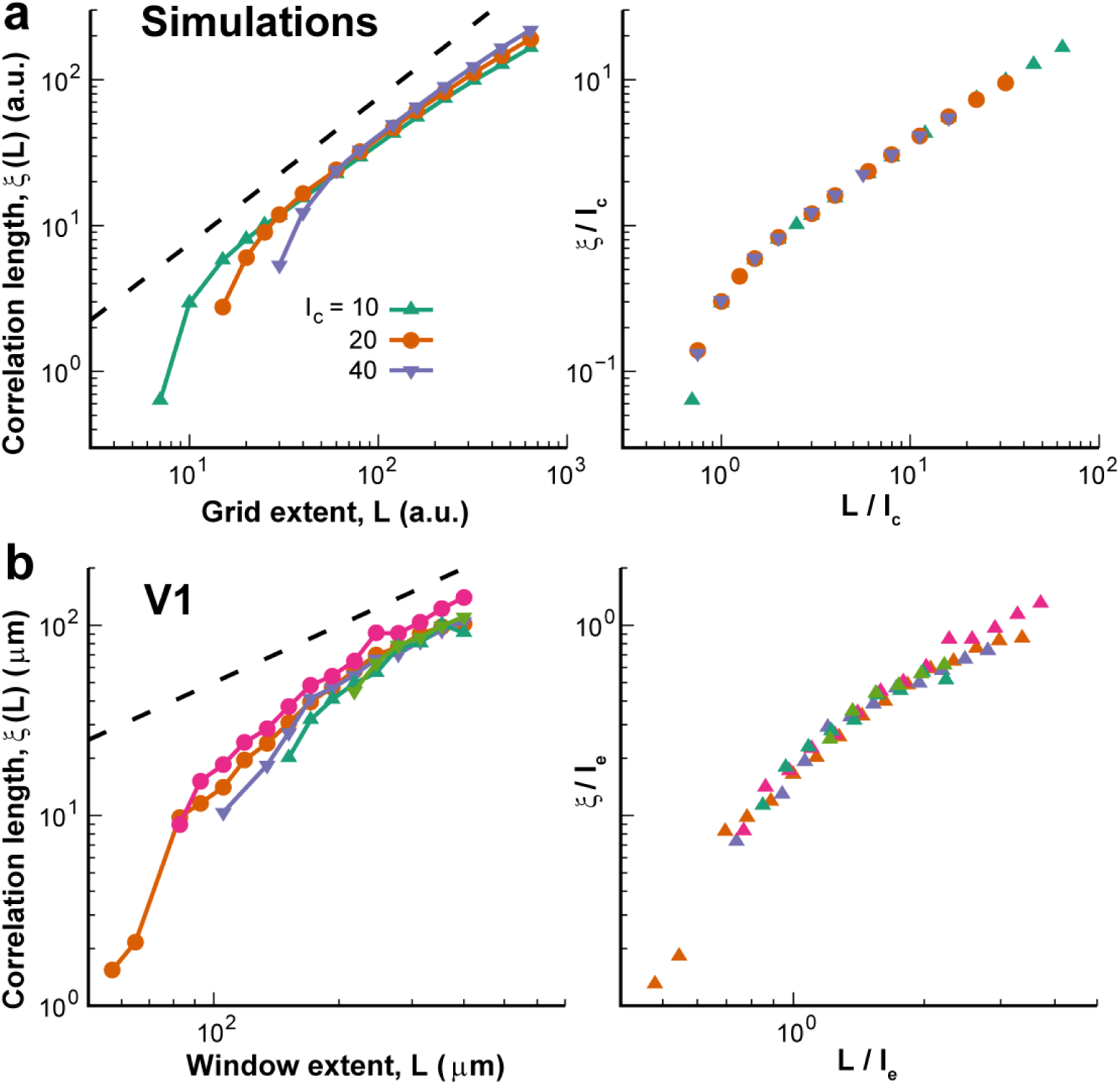
Interaction length in experimental data can be estimated from correlation length growth with observed window size. **a** *Left*: Correlation length ξ as a function of *L* for simulations at criticality for 3 different values of interaction length *I*_*c*_ (*color code*) in log-log coordinates. *Dashed line*: linear growth as a visual guide. *Right*: After rescaling grid extents and correlation lengths by the interaction length, all curves collapse into a single scaling function that highlights the change in the linear behavior at *L* = *I*_*c*_. In all cases, system size is 1000 x 1000 with connectivity *K* = 16. **b** Same as panel **a**, but for correlation lengths obtained from individual mice in V1. We estimated the point where the linear growth of the correlation length breaks for each mouse (see Materials and Methods) and used those values to rescale the curves as in panel **a** (*right*). The collapse obtained suggests that the estimated points (∼119 µm for mouse 1 – *orange*; ∼143 µm for mouse 2 – *purple*; ∼108 µm for mouse 3 – *pink*; ∼179 µm for mouse 4 – *dark green*; ∼180 µm for mouse 5 – *light green*) represent well the interaction length for these mice.

## Notes

### Competing Interest Statement

The authors have declared no competing interest.

